# SARS-CoV-2 ORF3b is a potent interferon antagonist whose activity is further increased by a naturally occurring elongation variant

**DOI:** 10.1101/2020.05.11.088179

**Authors:** Yoriyuki Konno, Izumi Kimura, Keiya Uriu, Masaya Fukushi, Takashi Irie, Yoshio Koyanagi, So Nakagawa, Kei Sato

**Author notes:** Correspondence (K.S.). These authors contributed equally (Yoriyuki Konno, Izumi Kimura).

## Abstract

One of the features distinguishing SARS-CoV-2 from its more pathogenic counterpart SARS-CoV is the presence of premature stop codons in its *ORF3b* gene. Here, we show that SARS-CoV-2 *ORF3b* is a potent interferon antagonist, suppressing the induction of type I interferon more efficiently than its SARS-CoV ortholog. Phylogenetic analyses and functional assays revealed that SARS-CoV-2-related viruses from bats and pangolins also encode truncated *ORF3b* gene products with strong anti-interferon activity. Furthermore, analyses of more than 15,000 SARS-CoV-2 sequences identified a natural variant, in which a longer *ORF3b* reading frame was reconstituted. This variant was isolated from two patients with severe disease and further increased the ability of ORF3b to suppress interferon induction. Thus, our findings not only help to explain the poor interferon response in COVID-19 patients, but also describe a possibility of the emergence of natural SARS-CoV-2 quasispecies with extended *ORF3b* that may exacerbate COVID-19 symptoms.

**Highlights:**

- ORF3b of SARS-CoV-2 and related bat and pangolin viruses is a potent IFN antagonist
- SARS-CoV-2 ORF3b suppresses IFN induction more efficiently than SARS-CoV ortholog
- The anti-IFN activity of ORF3b depends on the length of its C-terminus
- An ORF3b with increased IFN antagonism was isolated from two severe COVID-19 cases

## Introduction

In December 2019, an unusual outbreak of infectious pneumonia was reported in the city of Wuhan, Hubei, China. A few weeks later, a novel coronavirus (CoV) was identified as the causative agent and the disease was termed coronavirus disease 2019 (COVID-19) (Zhou et al., 2020). Since this novel virus is phylogenetically related to severe acute respiratory syndrome (SARS) CoV (SARS-CoV), it was named SARS-CoV-2. As of May 2020, SARS-CoV-2 causes an ongoing pandemic, with more than 4 million reported cases and more than 280,000 deaths worldwide (WHO, 2020).

SARS-CoV-2 infection may be asymptomatic or result in flu-like symptoms such as fever, cough and fatigue (Chen et al., 2020). In some cases, however, COVID-19 progresses to severe pneumonia and death (Guan et al., 2020; Hui et al., 2020; Li et al., 2020). Although it is still challenging to assess the morbidity rate of COVID-19, estimates range from 1.4 to 1.9% in China (Guan et al., 2020; Verity et al., 2020). This is substantially lower than the morbidity rate of SARS-CoV, which is about 9.6% (WHO, 2004). SARS-CoV, which frequently causes severe respiratory symptoms including fatal pneumonia, first emerged in Guangdong, China in 2002 and was stamped out in 2004 [reviewed in (Chan-Yeung and Xu, 2003; Weiss, 2020)]. Until then, 8,096 cases of SARS were reported in 29 countries and territories, and 774 people died (WHO, 2004). Thus, SARS-CoV is more virulent than SARS-CoV-2.

SARS-CoV-2 and SARS-CoV are phylogenetically closely related, both belonging to the family *Coronaviridae*, genus *Betacoronavirus* and subgenus *Sarbecovirus* (Lam et al., 2020; Zhou et al., 2020). Both viruses were transmitted from animals to humans. Thus, elucidating their zoonotic origin and phylogenetic history may help to understand genetic and phenotypic differences between SARS-CoV and SARS-CoV-2. Viruses closely related to SARS-CoV were detected in Chinese rufous horseshoe bats (*Rhinolophus sinicus*) (Lau et al., 2005; Li et al., 2005) and palm civets (*Paguma larvata*) (Wang et al., 2005). Subsequent surveillance studies have identified additional clades of SARS-CoV-related viruses in various bat species (mainly of the genus *Rhinolophus*) (Ge et al., 2013; He et al., 2014; Hu et al., 2017a; Lau et al., 2010; Lin et al., 2017; Tang et al., 2006; Wang et al., 2017; Wu et al., 2016; Yuan et al., 2010), suggesting that zoonotic coronavirus transmission from horseshoe bats to humans led to the emergence of SARS-CoV. Similarly, SARS-CoV-2-related viruses were identified in intermediate horseshoe bats (*Rhinolophus affinis*) (Zhou et al., 2020), the Malayan horseshoe bat (*Rhinolophus malayanus*) and Malayan pangolins (*Manis javanica*) (Lam et al., 2020; Xiao et al., 2020). Although it has been suggested that the SARS-CoV-2 outbreak has originated from cross-species coronavirus transmission from these mammals to humans, the exact origin remains to be determined (Andersen et al., 2020).

One prominent feature that distinguishes COVID-19 from SARS in terms of immune responses is the poor induction of a type I interferon (IFN-I) response by SARS-CoV-2 compared to SARS-CoV and influenza A virus (IAV) (Blanco-Melo et al., 2020; Hadjadj et al., 2020). Notably, impaired IFN-I responses are associated with COVID-19 disease (Hadjadj et al., 2020). However, the molecular mechanisms underlying the inefficient IFN-I responses in SARS-CoV-2 infection remain unclear. In this study, we therefore aimed to characterize the viral factor(s) determining immune activation upon SARS-CoV-2 infection. We particularly focused on differences in putative viral IFN-I antagonists and revealed that the *ORF3b* gene products of SARS-CoV-2 and SARS-CoV not only differ considerably in their length, but also in their ability to antagonize type I IFN. Furthermore, we demonstrate that the potent anti-IFN-I activity of SARS-CoV-2 ORF3b is also found in related viruses from bats and pangolins. Mutational analyses revealed that the length of the C-terminus determines the efficacy of IFN antagonism by ORF3b. Finally, we describe a natural SARS-CoV-2 variant with further increased ORF3b-mediated anti-IFN-I activity that emerged during the current COVID-19 pandemic.

## Results

### SARS-CoV-2 ORF3b is a potent IFN-I antagonist

To determine virological differences between SARS-CoV-2 and SARS-CoV, we set out to compare the sequences of diverse *Sarbecoviruses*. Consistent with recent reports (Lam et al., 2020; Zhou et al., 2020), *Sarbecoviruses* clustered into two groups, SARS-CoV-2-related and SARS-CoV-related viruses (**Figure 1A**). A comparison of individual viral open reading frames (ORFs) revealed that the length of ORF3b is clearly different between SARS-CoV-2 and SARS-CoV lineages, while the lengths of all remaining ORFs are relatively constant among *Sarbecoviruses* (**Figure 1B**). More specifically, the ORF3b sequences of SARS-CoV-2 and related viruses in bats and pangolins are only 22 amino acids (66 bp) long and therefore considerably shorter than those of their SARS-CoV orthologs (153.2 ± 0.47 amino acids on average).

**Figure 1.**
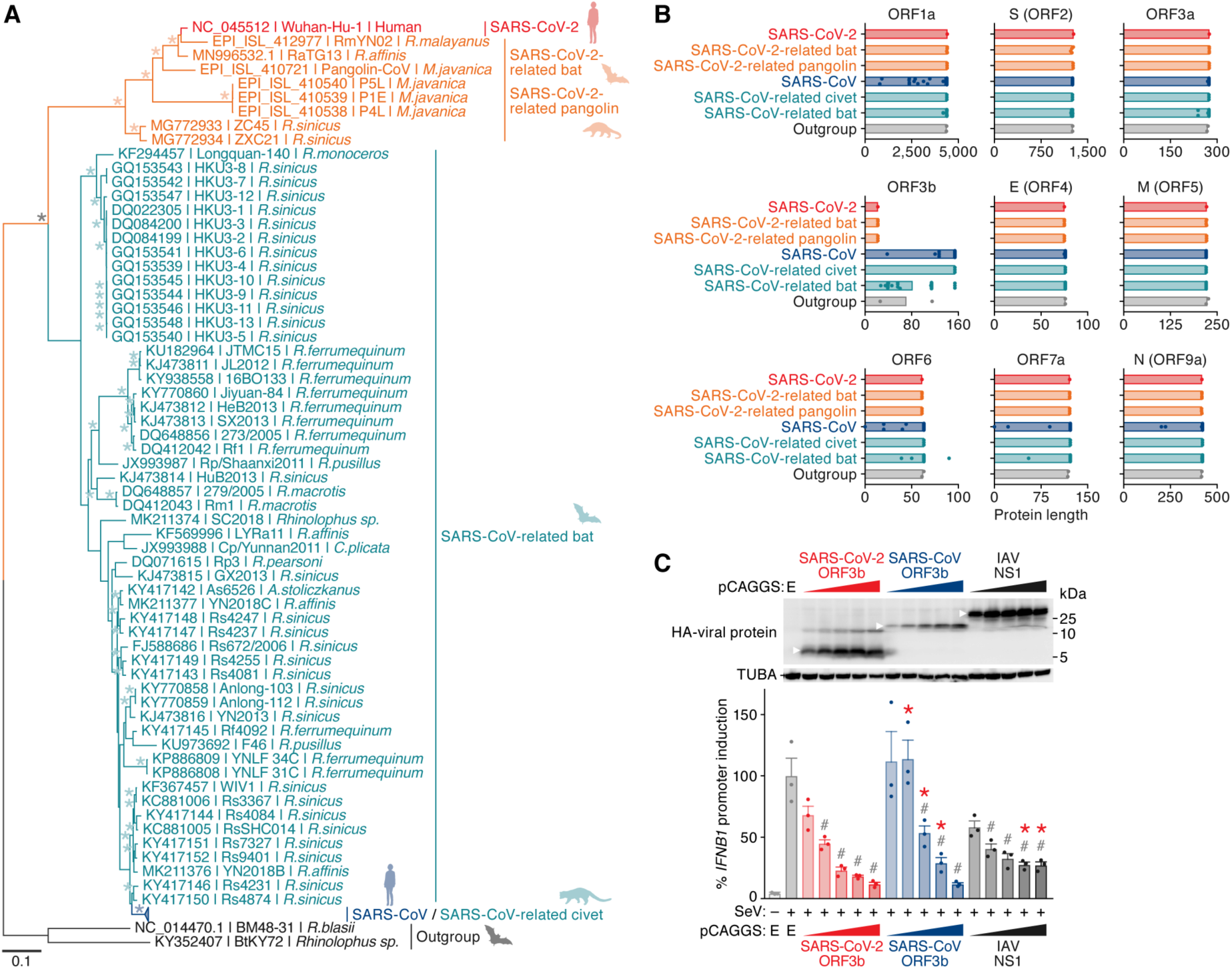
SARS-CoV-2 ORF3b is a potent IFN-I antagonist. **(A)** Maximum likelihood phylogenetic tree of full-length *Sarbecovirus* sequences. The full-length sequences (∼30,000 bp) of SARS-CoV-2 (Wuhan-Hu-1 as a representative), SARS-CoV-2-related viruses from bats (n=4) and pangolins (n=4), SARS-CoV (n=190), SARS-CoV-related viruses from civets (n=3) and bats (n=54), and outgroup viruses (n=2; BM48-31 and BtKY72) were analyzed. Accession number, strain name, and host of each virus are indicated for each branch. Note that the branches including SARS-CoV (n=190) and SARS-CoV-related viruses from civets (n=3) were collapsed for better visualization. The uncollapsed tree is shown in **Figure S1**, and the sequences used are summarized in **Table S1**. Asterisks indicate bootstrap values >95%. A scale bar indicates 0.1 nucleotide substitutions per site. NA, not applicable. **(B)** Comparison of the protein lengths of *Sarbecovirus* ORFs. The amino acid numbers of ORF1a, S (ORF2), ORF3a, ORF3b, E (ORF4), M (ORF5), ORF6, ORF7a, and N (ORF9a) of *Sarbecoviruses* are shown. The viral sequences used correspond to those in **A**. Bars indicate average values, and each dot represents one viral strain. ORFs with low similarity (e.g., ORF8 and ORF9b) were excluded from this analysis. **(C)** Potent anti-IFN-I activity of SARS-CoV-2 ORF3b. HEK293 cells were cotransfected with five different amounts of plasmids expressing HA-tagged SARS-CoV-2 ORF3b, SARS-CoV ORF3b, and IAV NS1 (50, 100, 200, 300, and 500 ng) and p125Luc, a plasmid encoding firefly luciferase under the control of the human *IFNB1* promoter (500 ng). 24 h post transfection, SeV was inoculated at MOI 10. 24 h post infection, the cells were harvested for Western blotting (top) and luciferase assay (bottom). For Western blotting, the input of cell lysate was normalized to TUBA, and one representative result out of X independent experiments is shown. The band of each viral protein is indicated by a white arrowhead. kDa, kilodalton. In the luciferase assay, the value of the SeV-infected empty vector-transfected cells was set to 100%. The average of three independent experiments with SEM is shown, and statistically significant differences (*P* < 0.05) compared to the SeV-infected empty vector-transfected cells (#) and the same amount of the SARS-CoV-2 ORF3b-transfected cells (*) are shown. E, empty vector. See also **Figure S1** and **Table S1**.

Previous studies on SARS-CoV and related viruses demonstrated that at least two accessory proteins, ORF3b and ORF6, as well as the nucleocapsid (N, also known as ORF9a) have the ability to inhibit IFN-I production (Frieman et al., 2007; Hu et al., 2017b; Kopecky-Bromberg et al., 2007; Zhou et al., 2012). Since the ORF3b length was remarkably different between SARS-CoV-2 and SARS-CoV (**Figure 1B**), we hypothesized that the antagonistic activity of ORF3b against IFN-I differs between these two viruses. To test this hypothesis, we monitored human *IFNB1* promoter activity in the presence of ORF3b of SARS-CoV-2 (Wuhan-Hu-1) and SARS-CoV (Tor2) using a luciferase reporter assay. The influenza A virus (IAV) non-structural protein 1 (NS1) served as positive control (Garcia-Sastre et al., 1998; Krug et al., 2003). As shown in **Figure 1C**, all three viral proteins dose-dependently suppressed the activation of the *IFNB1* promoter upon Sendai virus (SeV) infection. Notably, the antagonistic activity of SARS-CoV-2 ORF3b was slightly, but significantly higher than that of SARS-CoV ORF3b (**Figure 1C, bottom**). Thus, our data demonstrate that SARS-CoV-2 ORF3b is a potent inhibitor for human IFN-I activation, even though it only comprises 22 amino acids.

### SARS-CoV-2-related ORF3b proteins from bat and pangolin viruses suppress IFN-I activation on average more efficiently than their SARS-CoV counterparts

Since the lengths of ORF3b proteins in SARS-CoV-2-related viruses including those from bats and pangolins were on average shorter than those from SARS-CoV and related viruses (**Figure 1B**), we next investigated whether they were also generally more efficient in antagonizing IFN-I. A phylogenetic analysis of *Sarbecovirus ORF3b* genes showed that the evolutionary relationship of *Sarbecovirus ORF3b* genes was similar to that of the full-length viral genomes (**Figures 1A and 2A**). For our functional analyses, we generated expression plasmids for ORF3b from SARS-CoV-2-related viruses from bats (RmYN02, RaTG13 and ZXC21) and a pangolin (P4L), as well as SARS-CoV-related viruses from a civet (civet007) and bats (Rs7327, Rs4231, YN2013 and Rm1), representing different lengths of this protein (**Figure 2B**). As shown in **Figure 2C**, all four SARS-CoV-2-related ORF3b significantly suppressed human IFN-I activation. In contrast, only two SARS-CoV-related ORF3b proteins, Rs4231 and Rm1, exhibited anti-IFN-I activity at the concentrations tested (**Figure 2C**). Intriguingly, these two SARS-CoV-related ORF3b proteins are C-terminally truncated and shorter than ORF3b of SARS-CoV Tor2 (**Figure 2A**). These findings suggest that the C-terminal region (residues 115-154) may attenuate the anti-IFN-I activity of ORF3b. To test this hypothesis, we generated a C-terminally truncated derivatives of SARS-CoV (Tor2) ORF3b harboring a premature stop codon at position 135. The K135* mutant mimics the ORF3b of Rs4231 (**Figure 2B**). Reporter assays revealed that this derivative exhibits higher anti-IFN-I activity than wild-type (WT) SARS-CoV ORF3b (**Figure 2D**), demonstrating that the C-terminal region of SARS-CoV ORF3b indeed attenuates its anti-IFN-I activity.

**Figure 2.**
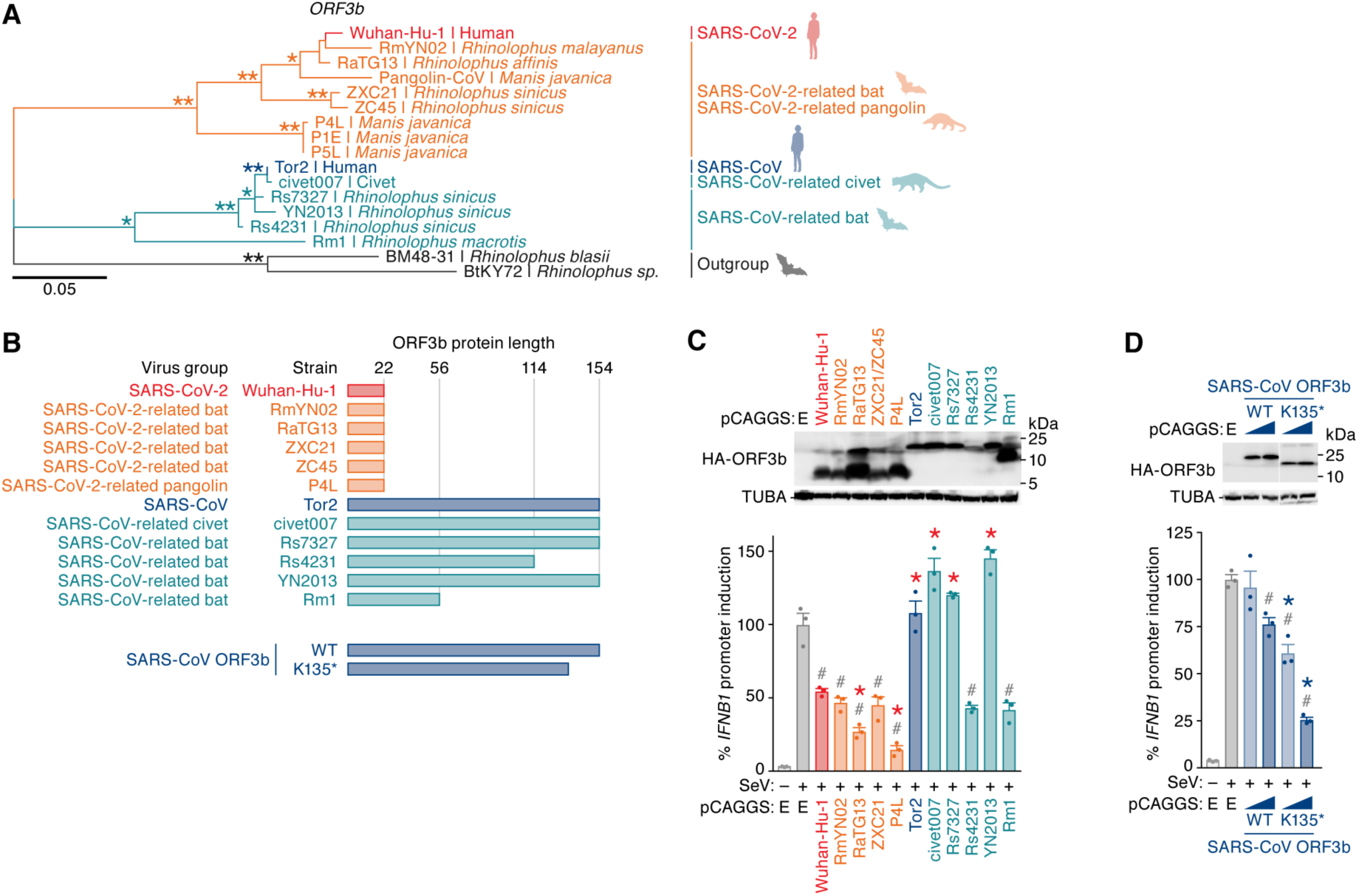
C-terminal truncations increase the IFN-antagonistic activity of ORF3b. **(A)** Maximum likelihood phylogenetic tree of *Sarbecovirus ORF3b*. The *ORF3b* sequences of SARS-CoV-2 (Wuhan-Hu-1), all reported SARS-CoV-2-related viruses from bats (n=4) and pangolins (n=4), representatives of SARS-CoV (Tor2), SARS-CoV-related viruses from civets (civet007) and bats (Rs7237, Rs4231, YN2013 and Rm1), and two outgroup viruses (BM48-31 and BtKY72) were analyzed. Strain name and host of each virus are indicated for each branch. Bootstrap value; *, >80%; **, >95%. **(B)** Illustration or protein lengths of all *Sarbecovirus* ORF3b isolates used in this study (top) and the protein lengths of SARS-CoV ORF3b mutants (bottom). **(C)** Anti-IFN-I activities of different *Sarbecovirus* ORF3b proteins. HEK293T cells were cotransfected with a plasmid expressing one of 11 HA-tagged *Sarbecovirus* ORF3b proteins (summarized in **B**; 100 ng) and p125Luc (500 ng). 24 h post transfection, SeV was inoculated at MOI 10. 24 h post infection, cells were harvested for Western blotting (top) and luciferase assay (bottom). Note that the amino acid sequences of ZXC21 and ZC45 are identical. **(D)** Anti-IFN-I activity of C-terminally truncated SARS-CoV ORF3b. HEK293T cells were cotransfected with two different amounts of plasmids expressing HA-tagged SARS-CoV ORF3b WT and K135* (50 and 100 ng) and p125Luc (500 ng). 24 h post transfection, SeV was inoculated at MOI 10. 24 h post infection, the cells were harvested for Western blotting (top) and luciferase assay (bottom). For Western blotting, the input of cell lysate was normalized to TUBA. One representative blot out of three independent experiments is shown. In the luciferase assay, the value of the SeV-infected empty vector-transfected cells was set to 100%. The average of three independent experiments with SEM is shown, and statistically significant differences (*P* < 0.05) compared to the SeV-infected empty vector-transfected cells (#) and the same amount of either SARS-CoV-2 ORF3b (Wuhan-Hu-1)-transfected cells (*, **C**) or SARS-CoV ORF3b WT-transfected cells (*, **D**) are shown. E, empty vector.

### A SARS-CoV *ORF3b*-like sequence is hidden in the SARS-CoV-2 genome

ORF3b of SARS-CoV-2 is shorter than its ortholog in SARS-CoV (**Figures 1B and 2A**). However, when closely inspecting the nucleotide sequences of these two viruses, we noticed that the SARS-CoV-2 nucleotide sequence downstream of the stop codon of *ORF3b* shows a high similarity to the SARS-CoV *ORF3b* gene (nucleotide similarity=79.5%; **Figure 3A**). In contrast to SARS-CoV *ORF3b*, however, SARS-CoV-2 harbors four premature stop codons that result in the expression of a drastically shortened ORF3b protein (**Figure 3A**). Similar patterns were observed in SARS-CoV-2-related viruses from bats and pangolins. Since the *ORF3b* length is closely associated with its anti-IFN-I activity (**Figures 2C and 2D**), we hypothesized that reversion of the premature stop codons in SARS-CoV-2 *ORF3b* affects its ability to inhibit human IFN-I. To address this possibility, we generated four SARS-CoV-2 ORF3b derivatives, 57*, 79*, 119* and 155*, lacking the respective premature stop codons (**Figure 3B, top**). As shown in **Figure 3C**, all four derivatives inhibited human IFN-I activation in dose-dependent manners. Consistent with the results obtained with SARS-CoV ORF3b mutants (**Figure 2D**), the 155* mutant, comprising the very C-terminal region (positions 119-154), was poorly expressed and exhibited relatively low anti-IFN-I activity (**Figure 3C**). Notably, however, we found that the extended ORF3b derivatives, particularly 57*, 79*, 119*, exhibited higher anti-IFN-I activity compared to WT ORF3b (**Figure 3C**). These findings confirm that the length of ORF3b determines its ability to suppress an IFN-I response. Furthermore, they show that the loss of authentic *ORF3b* stop codon during the current SARS-CoV-2 pandemic may result in the emergence of viral variants with enhanced IFN-I-antagonistic activity.

**Figure 3.**
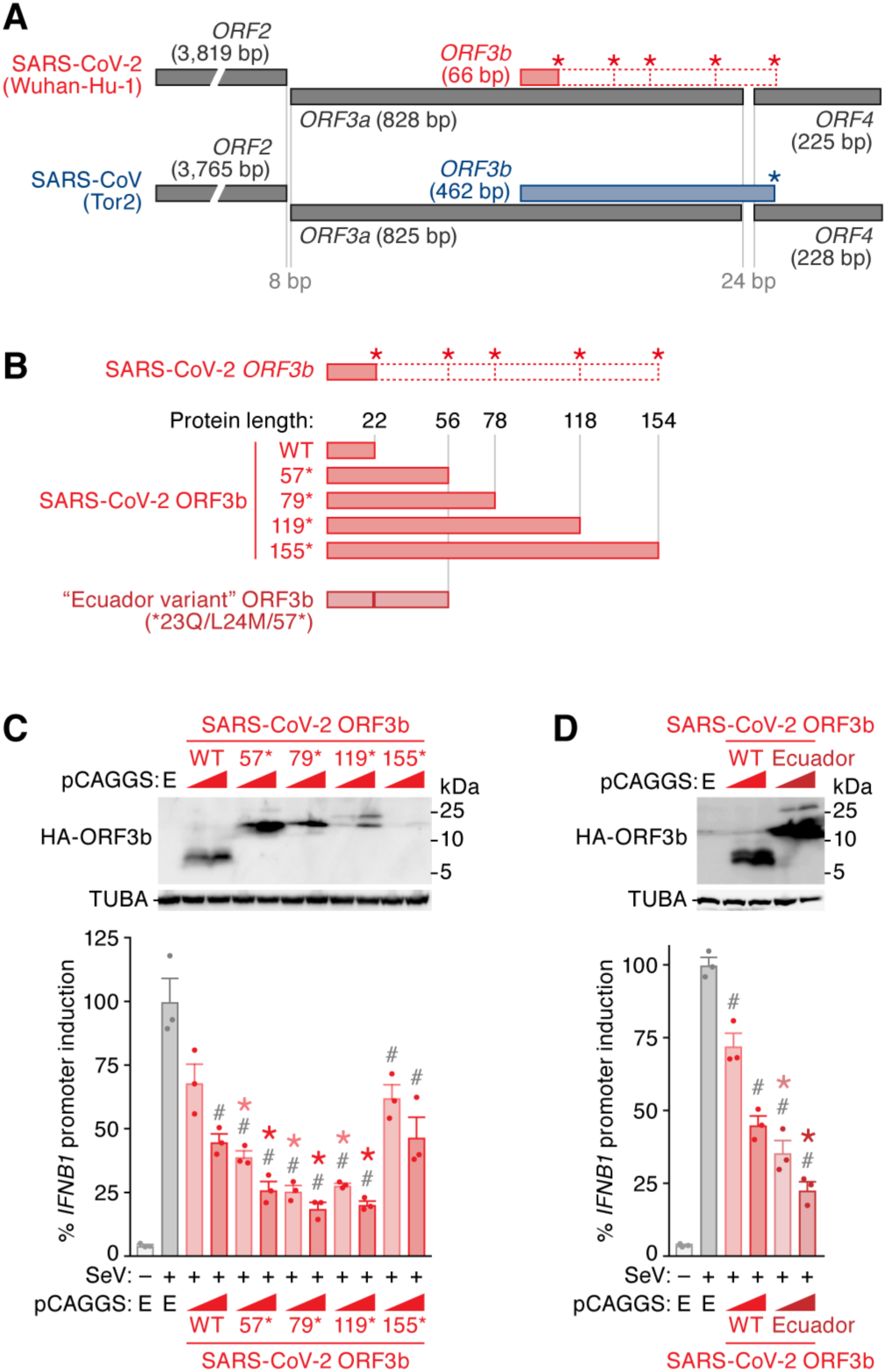
Enhanced anti-IFN-I upon reconstitution of the cryptic SARS-CoV-2 ORF3b. **(A)** Schemes illustrating the genomic regions encoding *ORF2, ORF3a, ORF3b* and *ORF4* of SARS-CoV-2 and SARS-CoV. Open squares with dotted red lines indicate a cryptic *ORF3b* reading frame in SARS-CoV-2 that is similar to SARS-CoV *ORF3b*. Asterisks indicate stop codons in the *ORF3b* frame. **(B)** SARS-CoV-2 ORF3b derivatives characterized in this study. (Top) WT SARS-CoV-2 ORF3b as well as four derivatives with mutated stop codons (57*, 79*, 119* and 155*) are shown. Asterisks indicate the stop codons in the original *ORF3b* frame. (Bottom) A natural ORF3b variant detected in two sequences deposited in GISAID (accession IDs: EPI_ISL_422564 and EPI_ISL_422565; herein designated an “Ecuador variant”) are shown. **(C)** Anti-IFN-I activity different SARS-CoV-2 ORF3b derivatives. HEK293T cells were cotransfected with two different amounts of plasmids expressing the indicated HA-tagged SARS-CoV-2 ORF3b derivatives (WT, 57*, 79*, 119* and 155*; 50 and 100 ng) and p125Luc (500 ng). 24 h post transfection, SeV was inoculated at MOI 10. 24 h post infection, the cells were harvested for Western blotting (top) and luciferase assay (bottom). **(D)** Enhanced anti-IFN-I activity of an “Ecuador variant” ORF3b. HEK293T cells were cotransfected with two different amounts of plasmids expressing HA-tagged “Ecuador variant” ORF3b or parental SARS-CoV-2 ORF3b (50 and 100 ng) and p125Luc (500 ng). 24 h post transfection, SeV was inoculated at MOI 10. 24 h post infection, the cells were harvested for Western blotting (top) and luciferase assay (bottom). For Western blotting, the input of cell lysate was normalized to TUBA. One representative blot out of three independent experiments is shown. A highly exposed blot visualizing the band of the 155* mutant is shown in **Figure S2**. kDa, kilodalton. In the luciferase assay, the value of the SeV-infected empty vector-transfected cells was set to 100%. The average of three independent experiments with SEM is shown, and statistically significant differences (*P* < 0.05) compared to the SeV-infected empty vector-transfected cells (#) and the same amount of the SARS-CoV-2 ORF3b WT-transfected cells (*) are shown. E, empty vector. See also **Figure S2**.

### Characterization of a natural SARS-CoV-2 ORF3b variant with enhanced anti-IFN-I activity

By screening >15,000 viral sequences deposited in GISAID (https://www.gisaid.org; as of 22 April, 2020) using the CoV-GLUE webtool (http://cov-glue.cvr.gla.ac.uk), we detected two viral sequences (accession IDs: EPI_ISL_422564 and EPI_ISL_422565), in which the *ORF3b* gene was extended due to the loss of the first premature stop codon (*23Q) (**Figure 3B, bottom**). The similarity of the full-length sequences of these two viruses, which were collected from COVID-19 patients in Ecuador, were >99.6%, and the *ORF3b* sequences were identical. Apart from the *23Q mutation, the “Ecuador variant” also harbored an L24M change compared to the SARS-CoV *ORF3b*-like sequence in SARS-CoV-2 [Wuhan-Hu-1 (accession no. NC_045512.2), nucleotides 25814-26281] (**Figure 3B, bottom**). IFNβ reporter assays revealed that the “Ecuador variant” ORF3b exhibits significantly higher anti-IFN-I activity than the parental SARS-CoV-2 ORF3b (**Figure 3D**). These findings show that a naturally occurring SARS-CoV-2 variant, expressing an elongated ORF3b protein with enhanced anti-IFN activity, has already emerged during the current SARS-CoV-2 pandemic.

## Discussion

Here, we demonstrate that SARS-CoV-2 ORF3b is a potent antagonist of human IFN-I activation. On average, ORF3b proteins from SARS-CoV-2 and related bat and pangolin viruses were more active than their SARS-CoV counterparts. Notably, a recent study has revealed that the antibodies recognizing ORF3b are highly detectable during the early phase of SARS-CoV-2 infection (Hachim et al., 2020), which suggests that ORF3b is one of the viral proteins dominantly expressed in the COVID-19 patients during acute infection. Therefore, our findings may help to explain the inefficient and delayed IFN-I responses in SARS-CoV-2-infected cells as well as COVID-19 patients (Blanco-Melo et al., 2020). Moreover, a recent study showed that impaired IFN-I responses as well as reduced IFN-stimulated gene expression are associated with severe COVID-19 disease (Hadjadj et al., 2020). This suggests that imbalanced IFN-I responses against SARS-CoV-2 infection may determine its pathogenicity and explain differences compared to SARS-CoV. Thus, it is tempting to speculate that atypical symptoms and poor IFN-I responses in SARS-CoV-2 infection may be attributed to the potent anti IFN-I activity of its ORF3b.

Like SARS-CoV-2 *ORF3b*, its orthologs in SARS-CoV-2-related viruses from bats and pangolins efficiently antagonize IFN-I and are generally truncated due to the presence of several premature stop codons. In contrast, the anti-IFN activity of ORF3b proteins encoded by some SARS-CoV-related viruses is attenuated, most likely due to an elongated C-terminus. We hypothesized that the *ORF3b* length variation in SARS-CoV-like viruses may be the result of recombination events. In line with this, *Sarbecoviruses* seem to easily recombine with each other (Andersen et al., 2020; Lam et al., 2020; Zhou et al., 2020), and some horseshoe bat species such as *Rhinolophus affinis* and *Rhinolophus sinicus* are known to harbor both SARS-CoV-2- and SARS-CoV-related viruses (Andersen et al., 2020; Zhou et al., 2020). Nevertheless, the phylogenetic topologies of the full-length viral genome and the *ORF3b* gene are similar, and we found no evidence for recombination of *ORF3b* between the lineages of SARS-CoV-2 and SARS-CoV. Notably, phenotypic differences in the ability of ORF3b to suppress IFN-I responses may also be associated with the likelihood of successful zoonotic transmission of *Sarbecoviruses* to humans since many IFN-stimulated genes are antagonized in a species-specific manner. While more than 50 SARS-CoV-related viruses were isolated from bats (Ge et al., 2013; He et al., 2014; Hu et al., 2017a; Lau et al., 2010; Lau et al., 2005; Li et al., 2005; Lin et al., 2017; Tang et al., 2006; Wang et al., 2017; Wu et al., 2016; Yuan et al., 2010), only eight viral sequences belonging to the SARS-CoV-2 lineage were detected so far (Andersen et al., 2020; Lam et al., 2020; Xiao et al., 2020; Zhou et al., 2020). Thus, further investigations are needed to elucidate the dynamics of cross-species transmission events of *Sarbecoviruses* and the evolution of the *ORF3b* gene.

We further show that a SARS-CoV *ORF3b*-like sequence is still present in the SARS-CoV-2 genome, but is interrupted by premature stop codons. We demonstrate that a partial extension of SARS-CoV-2 *ORF3b* by reverting stop codons increases its inhibitory activity against human IFN-I. Full reversion of all stop codons, however, resulted in an ORF3b protein with poor anti-IFN activity. This is in line with the phenotypic difference between SARS-CoV-2 and SARS-CoV ORF3b proteins and suggests that the very C-terminus of ORF3b impairs its immune evasion activity.

Intriguingly, we also identified a naturally occurring SARS-CoV-2 *ORF3b* variant that expresses an elongated protein due to the loss of the first premature stop codon. This variant suppresses IFN-I even more efficiently than ORF3b of the SARS-CoV-2 reference strain. In agreement with an association of IFN suppression with disease severity (Hadjadj et al., 2020), the two patients in Ecuador harboring SARS-CoV-2 with the extended *ORF3b* variant were critically ill; one (accession ID: EPI_ISL_422564) was treated in an intensive care unit and the other (accession ID: EPI_ISL_422565) died of COVID-19 (unpublished information from Dr. Paúl Cárdenas, Universidad San Francisco de Quito, Quito, Ecuador). There are no direct evidence indicating that the viruses detected in these two COVID-19 patients in Ecuador are highly pathogenic. However, based on our experimental results, it might be plausible to assume that naturally occurring length variants of *ORF3b* may occur due to the loss of premature stop codons and potentially contribute to the emergence of highly pathogenic SARS-CoV-2 quasispecies. Our findings can be a clue to monitor the possibility of the emergence of highly pathogenic viruses during current SARS-CoV-2 pandemic.

## Materials and Methods

### Cell Culture

HEK293 cells (a human embryonic kidney cell line; ATCC CRL-1573) were maintained in Dulbecco’s modified Eagle’s medium (Sigma-Aldrich) containing fetal calf serum and antibiotics.

### Viral Genomes and Phylogenetic Analyses

All viral genome sequences used in this study and the respective GenBank or GISAID (https://www.gisaid.org) accession numbers are summarized in **Table S1**. We first aligned the viral genomes using the L-INS-i program of MAFFT version 7.453 (Katoh and Standley, 2013). Based on the multiple sequence alignment and the gene annotation of SARS-CoV, we extracted the region of the *ORF3b* gene. We then constructed phylogenetic trees using the full-length genomes (**Figures 1A and S1**) and *ORF3b* gene (**Figure 2A**). We generated a maximum likelihood based phylogenetic tree using RAxML-NG (Kozlov et al., 2019) with a General Time Reversible model of nucleotide substitution with invariant sites and gamma distributed rate variation among sites. We visualized the tree using a FigTree software (http://tree.bio.ed.ac.uk/software/figtree).

### Plasmid Construction

To construct the expression plasmids for HA-tagged *Sarbecovirus* ORF3b and IAV A/Puerto Rico/8/34 (H1N1 PR8; accession no. EF467817.1) NS1, pCAGGS (Niwa et al., 1991) was used as a backbone. The HA-tagged ORF of each gene (the accession numbers and sequences are listed in **Table S2**) and the cryptic SARS-CoV *ORF3b-*like sequence in SARS-CoV-2 [Wuhan-Hu-1 (accession no. NC_045512.2), nucleotides 25814-26281) was synthesized by a gene synthesis service (Fasmac). The ORF3b derivatives were generated by PCR using PrimeSTAR GXL DNA polymerase (Takara), the synthesized ORFs as templates, and the primers listed in **Table S3**. The HA-tagged “Ecuador variant” ORF3b (accession IDs: EPI_ISL_422564 and EPI_ISL_422565, which corresponds to the S23Q/L24M mutant of SARS-CoV-2 Wuhan-Hu-1 ORF3b *57) was generated by overlap extension PCR by using PrimeSTAR GXL DNA polymerase (Takara), the SARS-CoV-2 ORF3b 155* as the template, and the primers listed in **Table S3**. The obtained DNA fragments were inserted into pCAGGS via EcoRI-BglII or XhoI-BglII. Nucleotide sequences were determined by a DNA sequencing service (Fasmac), and the sequence data were analyzed by Sequencher v5.1 software (Gene Codes Corporation).

### Transfection, SeV Infection and Reporter Assay

HEK293 cells were transfected using PEI Max (Polysciences) according to the manufacturer’s protocol. For luciferase reporter assay, cells were cotransfected with 500 ng of p125Luc (expressing firefly luciferase driven by human *IFNB1* promoter; kindly provided by Dr. Takashi Fujita) (Fujita et al., 1993) and the pCAGGS-based HA-tagged expression plasmid (the amounts are indicated in the figure legends). At 24 h posttransfection, SeV (strain Cantell, clone cCdi; accession no. AB855654) (Yoshida et al., 2018) was inoculated into the transfected cells at multiplicity of infection (MOI) 10. The luciferase reporter assay was performed as described (Kobayashi et al., 2014; Konno et al., 2018; Ueda et al., 2017). Briefly, 50 μl of cell lysate was applied to a 96-well plate (Nunc), and the firefly luciferase activity was measured using a BrillianStar-LT assay system (Toyo-b-net), and the input for the luciferase assay was normalized by using a CellTiter-Glo 2.0 assay kit (Promega) following the manufacturers’ instructions. For this assay, a 2030 ARVO X multilabel counter instrument (PerkinElmer) was used.

### Western Blotting

Western blotting was performed as described (Kobayashi et al., 2014; Konno et al., 2018; Nakano et al., 2017; Yamada et al., 2018) using an HRP-conjugated anti-HA rat monoclonal antibody (clone 3F10; Roche) and an anti-alpha-tubulin (TUBA) mouse monoclonal antibody (clone DM1A; Sigma-Aldrich). Transfected cells were lysed with RIPA buffer (25 mM HEPES [pH 7.4], 50 mM NaCl, 1 mM MgCl_2_, 50 μM ZnCl_2_, 10% glycerol, 1% Triton X-100) containing a protease inhibitor cocktail (Roche).

### CoV-GLUE

To survey the *ORF3b* derivatives in pandemic SARS-CoV-2 sequences, we used the 15,605 viral sequences deposited in GISAID (https://www.gisaid.org) (accessed 22 April, 2020). The screening was performed using the CoV-GLUE platform (http://cov-glue.cvr.gla.ac.uk) developed by MRC-University of Glasgow Centre for Virus Research, Scotland, UK (accessed 22 April, 2020). Using CoV-GLUE, we detected the two SARS-CoV-2 sequences (accession IDs: EPI_ISL_422564 and EPI_ISL_422565, collected in Ecuador) possessing the V163T/T164N substitutions in ORF3a, which correspond to the *23Q/L24M/57* substitutions in ORF3b.

### Statistical Analysis

Data analyses were performed using Prism 7 (GraphPad Software). The data are presented as averages ± SEM. Statistically significant differences were determined by Student’s *t* test. Statistical details can be found directly in the figures or in the corresponding figure legends.

## Author Contributions

Y.Konno, I.K., K.U., and T.I. performed the experiments.

S.N. performed molecular phylogenetic analysis.

T.I. and Y.Koyanagi prepared reagents.

Y.Konno, I.K., T.I. and K.S. interpreted the results.

K.S. designed the experiments and wrote the manuscript. All authors reviewed and proofread the manuscript.

## Acknowledgments

We would like to thank all laboratory members in Division of Systems Virology, Institute of Medical Science, the University of Tokyo, Japan, and all the authors who have kindly deposited and shared genome data on GISAID. We also thank Naoko Misawa, Masayuki Horie and Keizo Tomonaga (Institute for Life and Medical Sciences, Kyoto University, Japan) and Ryoko Kawabata (Institute of Biomedical and Health Sciences, Hiroshima University, Japan) for generous supports, Daniel Sauter (Ulm University, Germany) for providing critical comments and suggestions for this study, Ken Maeda (National Institute of Infectious Diseases, Japan) for providing BKT1 cells, and Takashi Fujita (Institute for Life and Medical Sciences, Kyoto University, Japan) for providing p125Luc. We would like to appreciate Paúl Cárdenas (Universidad San Francisco de Quito, Ecuador) for providing the clinical information of the two COVID-19 patients in Ecuador. We thank Kotubu Misawa for dedicated support.

This study was supported in part by AMED Research Program on Emerging and Re-emerging Infectious Diseases 20fk0108146h0001 (to K.S.); AMED Research Program on HIV/AIDS 19fk0410019h0003 (to Y.Koyanagi and K.S.) and 20fk0410014h0003 (to Y.Koyanagi and K.S.); KAKENHI Grant-in-Aid for Scientific Research B 18H02662 (to Y.S. and K.S.), KAKENHI Grant-in-Aid for Scientific Research on Innovative Areas 16H06429 (to S.N., T.I., and K.S.), 16K21723 (to S.N., T.I., and K.S.), 17H05823 (to S.N.), 17H05813 (to K.S.), 19H04837 (to T.I.), 19H04843 (to S.N.) and 19H04826 (to K.S.), and Fund for the Promotion of Joint International Research (Fostering Joint International Research) 18KK0447 (to K.S.); JSPS Research Fellow DC1 19J22914 (to Y.Konno), DC1 19J20488 (to I.K.); Takeda Science Foundation (to K.S.); ONO Medical Research Foundation (to K.S.); Ichiro Kanehara Foundation (to K.S.); Lotte Foundation (to K.S.); Mochida Memorial Foundation for Medical and Pharmaceutical Research (to K.S.); Daiichi Sankyo Foundation of Life Science (to K.S.); Sumitomo Foundation (to K.S.); Uehara Foundation (to K.S.); Joint Research Project of the Institute of Medical Science, the University of Tokyo (to Y.Koyanagi); Joint Usage/Research Center program of Institute for Frontier Life and Medical Sciences, Kyoto University (to K.S.); and JSPS Core-to-Core program (A. Advanced Research Networks) (to Y.Koyanagi and K.S.).

## References

Andersen, K.G., Rambaut, A., Lipkin, W.I., Holmes, E.C., and Garry, R.F. (2020). The proximal origin of SARS-CoV-2. Nat Med 26, 450–452.

Blanco-Melo, D., Nilsson-Payant, B.E., Liu, W.-C., Uhl, S., Hoagland, D., Møller, R., Jordan, T.X., Oishi, K., Panis, M., Sachs, D., et al. (2020). Imbalanced host response to SARS-CoV-2 drives development of COVID-19. Cell in press.

Chan-Yeung, M., and Xu, R.H. (2003). SARS: epidemiology. Respirology 8 Suppl, S9–14.

Chen, N., Zhou, M., Dong, X., Qu, J., Gong, F., Han, Y., Qiu, Y., Wang, J., Liu, Y., Wei, Y., et al. (2020). Epidemiological and clinical characteristics of 99 cases of 2019 novel coronavirus pneumonia in Wuhan, China: a descriptive study. Lancet 395, 507–513.

Frieman, M., Yount, B., Heise, M., Kopecky-Bromberg, S.A., Palese, P., and Baric, R.S. (2007). Severe acute respiratory syndrome coronavirus ORF6 antagonizes STAT1 function by sequestering nuclear import factors on the rough endoplasmic reticulum/Golgi membrane. J Virol 81, 9812–9824.

Fujita, T., Nolan, G.P., Liou, H.C., Scott, M.L., and Baltimore, D. (1993). The candidate proto-oncogene bcl-3 encodes a transcriptional coactivator that activates through NF-kappa B p50 homodimers. Genes Dev 7, 1354–1363.

Garcia-Sastre, A., Egorov, A., Matassov, D., Brandt, S., Levy, D.E., Durbin, J.E., Palese, P., and Muster, T. (1998). Influenza A virus lacking the NS1 gene replicates in interferon-deficient systems. Virology 252, 324–330.

Ge, X.Y., Li, J.L., Yang, X.L., Chmura, A.A., Zhu, G., Epstein, J.H., Mazet, J.K., Hu, B., Zhang, W., Peng, C., et al. (2013). Isolation and characterization of a bat SARS-like coronavirus that uses the ACE2 receptor. Nature 503, 535–538.

Guan, W.J., Ni, Z.Y., Hu, Y., Liang, W.H., Ou, C.Q., He, J.X., Liu, L., Shan, H., Lei, C.L., Hui, D.S.C., et al. (2020). Clinical characteristics of coronavirus disease 2019 in China. N Engl J Med 382, 1708–1720.

Hachim, A., Kavian, N., Cohen, C.A., Chin, A.W., Chu, D.K., Mok, C.K.P., Tsang, O.T., Yeung, Y.C., Perera, R.A., Poon, L.L., et al. (2020). Beyond the Spike: identification of viral targets of the antibody response to SARS-CoV-2 in COVID-19 patients. MedRxiv, 20085670.

Hadjadj, J., Yatim, N., Barnabei, L., Corneau, A., Boussier, J., Pere, H., Charbit, B., Bondet, V., Chenevier-Gobeaux, C., Breillat, P., et al. (2020). Impaired type I interferon activity and exacerbated inflammatory responses in severe Covid-19 patients. MedRxiv, 20068015.

He, B., Zhang, Y., Xu, L., Yang, W., Yang, F., Feng, Y., Xia, L., Zhou, J., Zhen, W., Feng, Y., et al. (2014). Identification of diverse alphacoronaviruses and genomic characterization of a novel severe acute respiratory syndrome-like coronavirus from bats in China. J Virol 88, 7070–7082.

Hu, B., Zeng, L.P., Yang, X.L., Ge, X.Y., Zhang, W., Li, B., Xie, J.Z., Shen, X.R., Zhang, Y.Z., Wang, N., et al. (2017a). Discovery of a rich gene pool of bat SARS-related coronaviruses provides new insights into the origin of SARS coronavirus. PLoS Pathog 13, e1006698.

Hu, Y., Li, W., Gao, T., Cui, Y., Jin, Y., Li, P., Ma, Q., Liu, X., and Cao, C. (2017b). The severe acute respiratory syndrome coronavirus nucleocapsid inhibits type I interferon production by interfering with TRIM25-mediated RIG-I ubiquitination. J Virol 91.

Hui, D.S., E, I.A., Madani, T.A., Ntoumi, F., Kock, R., Dar, O., Ippolito, G., McHugh, T.D., Memish, Z.A., Drosten, C., et al. (2020). The continuing 2019-nCoV epidemic threat of novel coronaviruses to global health - The latest 2019 novel coronavirus outbreak in Wuhan, China. Int J Infect Dis 91, 264–266.

Katoh, K., and Standley, D.M. (2013). MAFFT multiple sequence alignment software version 7: improvements in performance and usability. Mol Biol Evol 30, 772–780.

Kobayashi, T., Takeuchi, J.S., Ren, F., Matsuda, K., Sato, K., Kimura, Y., Misawa, N., Yoshikawa, R., Nakano, Y., Yamada, E., et al. (2014). Characterization of red-capped mangabey tetherin: implication for the co-evolution of primates and their lentiviruses. Sci Rep 4, 5529.

Konno, Y., Nagaoka, S., Kimura, I., Takahashi Ueda, M., Kumata, R., Ito, J., Nakagawa, S., Kobayashi, T., Koyanagi, Y., and Sato, K. (2018). A naturally occurring feline APOBEC3 variant that loses anti-lentiviral activity by lacking two amino acid residues. J Gen Virol 99, 704–709.

Kopecky-Bromberg, S.A., Martinez-Sobrido, L., Frieman, M., Baric, R.A., and Palese, P. (2007). Severe acute respiratory syndrome coronavirus open reading frame (ORF) 3b, ORF 6, and nucleocapsid proteins function as interferon antagonists. J Virol 81, 548–557.

Kozlov, A.M., Darriba, D., Flouri, T., Morel, B., and Stamatakis, A. (2019). RAxML-NG: a fast, scalable and user-friendly tool for maximum likelihood phylogenetic inference. Bioinformatics 35, 4453–4455.

Krug, R.M., Yuan, W., Noah, D.L., and Latham, A.G. (2003). Intracellular warfare between human influenza viruses and human cells: the roles of the viral NS1 protein. Virology 309, 181–189.

Lam, T.T., Shum, M.H., Zhu, H.C., Tong, Y.G., Ni, X.B., Liao, Y.S., Wei, W., Cheung, W.Y., Li, W.J., Li, L.F., et al. (2020). Identifying SARS-CoV-2 related coronaviruses in Malayan pangolins. Nature.

Lau, S.K., Li, K.S., Huang, Y., Shek, C.T., Tse, H., Wang, M., Choi, G.K., Xu, H., Lam, C.S., Guo, R., et al. (2010). Ecoepidemiology and complete genome comparison of different strains of severe acute respiratory syndrome-related Rhinolophus bat coronavirus in China reveal bats as a reservoir for acute, self-limiting infection that allows recombination events. J Virol 84, 2808–2819.

Lau, S.K., Woo, P.C., Li, K.S., Huang, Y., Tsoi, H.W., Wong, B.H., Wong, S.S., Leung, S.Y., Chan, K.H., and Yuen, K.Y. (2005). Severe acute respiratory syndrome coronavirus-like virus in Chinese horseshoe bats. Proc Natl Acad Sci U S A 102, 14040–14045.

Li, Q., Guan, X., Wu, P., Wang, X., Zhou, L., Tong, Y., Ren, R., Leung, K.S.M., Lau, E.H.Y., Wong, J.Y., et al. (2020). Early transmission dynamics in Wuhan, China, of novel coronavirus-infected pneumonia. N Engl J Med 382, 1199–1207.

Li, W., Shi, Z., Yu, M., Ren, W., Smith, C., Epstein, J.H., Wang, H., Crameri, G., Hu, Z., Zhang, H., et al. (2005). Bats are natural reservoirs of SARS-like coronaviruses. Science 310, 676–679.

Lin, X.D., Wang, W., Hao, Z.Y., Wang, Z.X., Guo, W.P., Guan, X.Q., Wang, M.R., Wang, H.W., Zhou, R.H., Li, M.H., et al. (2017). Extensive diversity of coronaviruses in bats from China. Virology 507, 1–10.

Nakano, Y., Misawa, N., Juarez-Fernandez, G., Moriwaki, M., Nakaoka, S., Funo, T., Yamada, E., Soper, A., Yoshikawa, R., Ebrahimi, D., et al. (2017). Correction: HIV-1 competition experiments in humanized mice show that APOBEC3H imposes selective pressure and promotes virus adaptation. PLoS Pathog 13, e1006606.

Niwa, H., Yamamura, K., and Miyazaki, J. (1991). Efficient selection for high-expression transfectants with a novel eukaryotic vector. Gene 108, 193–199.

Tang, X.C., Zhang, J.X., Zhang, S.Y., Wang, P., Fan, X.H., Li, L.F., Li, G., Dong, B.Q., Liu, W., Cheung, C.L., et al. (2006). Prevalence and genetic diversity of coronaviruses in bats from China. J Virol 80, 7481–7490.

Ueda, M.T., Kurosaki, Y., Izumi, T., Nakano, Y., Oloniniyi, O.K., Yasuda, J., Koyanagi, Y., Sato, K., and Nakagawa, S. (2017). Functional mutations in spike glycoprotein of Zaire ebolavirus associated with an increase in infection efficiency. Genes Cells 22, 148–159.

Verity, R., Okell, L.C., Dorigatti, I., Winskill, P., Whittaker, C., Imai, N., Cuomo-Dannenburg, G., Thompson, H., Walker, P.G.T., Fu, H., et al. (2020). Estimates of the severity of coronavirus disease 2019: a model-based analysis. Lancet Infect Dis.

Wang, L., Fu, S., Cao, Y., Zhang, H., Feng, Y., Yang, W., Nie, K., Ma, X., and Liang, G. (2017). Discovery and genetic analysis of novel coronaviruses in least horseshoe bats in southwestern China. Emerg Microbes Infect 6, e14.

Wang, M., Yan, M., Xu, H., Liang, W., Kan, B., Zheng, B., Chen, H., Zheng, H., Xu, Y., Zhang, E., et al. (2005). SARS-CoV infection in a restaurant from palm civet. Emerg Infect Dis 11, 1860–1865.

Weiss, S.R. (2020). Forty years with coronaviruses. J Exp Med 217.

WHO (2004). “Summary of probable SARS cases with onset of illness from 1 November 2002 to 31 July 2003”. https://www.who.int/csr/sars/country/table2004_04_21/en/.

WHO (2020). “Coronavirus disease 2019”. https://www.who.int/emergencies/diseases/novel-coronavirus-2019.

Wu, Z., Yang, L., Ren, X., Zhang, J., Yang, F., Zhang, S., and Jin, Q. (2016). ORF8-related genetic evidence for Chinese horseshoe bats as the source of human severe acute respiratory syndrome coronavirus. J Infect Dis 213, 579–583.

Xiao, K., Zhai, J., Feng, Y., Zhou, N., Zhang, X., Zou, J.-J., Li, N., Guo, Y., Li, X., Shen, X., et al. (2020). Isolation of SARS-CoV-2-related coronavirus from Malayan pangolins. Nature in press.

Yamada, E., Nakaoka, S., Klein, L., Reith, E., Langer, S., Hopfensperger, K., Iwami, S., Schreiber, G., Kirchhoff, F., Koyanagi, Y., et al. (2018). Human-specific adaptations in Vpu conferring anti-tetherin activity are critical for efficient early HIV-1 replication in vivo. Cell Host Microbe 23, 110–120 e117.

Yoshida, A., Kawabata, R., Honda, T., Sakai, K., Ami, Y., Sakaguchi, T., and Irie, T. (2018). A Single Amino Acid Substitution within the Paramyxovirus Sendai Virus Nucleoprotein Is a Critical Determinant for Production of Interferon-Beta-Inducing Copyback-Type Defective Interfering Genomes. J Virol 92.

Yuan, J., Hon, C.C., Li, Y., Wang, D., Xu, G., Zhang, H., Zhou, P., Poon, L.L., Lam, T.T., Leung, F.C., et al. (2010). Intraspecies diversity of SARS-like coronaviruses in Rhinolophus sinicus and its implications for the origin of SARS coronaviruses in humans. J Gen Virol 91, 1058–1062.

Zhou, P., Li, H., Wang, H., Wang, L.F., and Shi, Z. (2012). Bat severe acute respiratory syndrome-like coronavirus ORF3b homologues display different interferon antagonist activities. J Gen Virol 93, 275–281.

Zhou, P., Yang, X.L., Wang, X.G., Hu, B., Zhang, L., Zhang, W., Si, H.R., Zhu, Y., Li, B., Huang, C.L., et al. (2020). A pneumonia outbreak associated with a new coronavirus of probable bat origin. Nature 579, 270–273.

