## Supplementary figures S1 and S2 for "SARS-CoV-2 ORF3b is a potent interferon antagonist whose activity is further increased by a naturally occurring elongation variant"

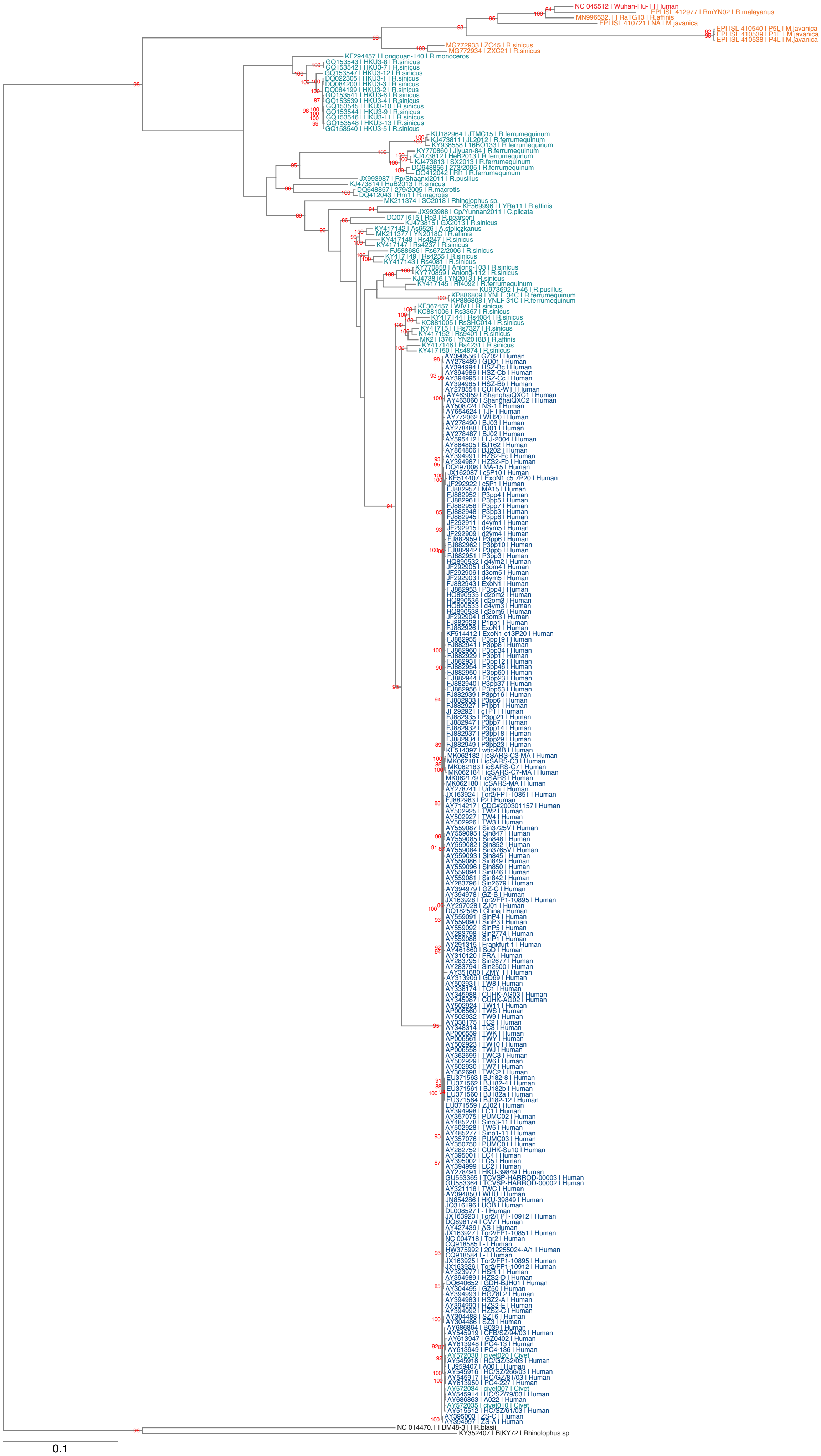
NC 045512 | Wuhan-Hu-1 | Human
EPI ISL 412977 | RmYN02 | R.malayanus
MN996532.1 | RaTG13 | R.affinis
EPI ISL 410721 | NA | M.javanica
EPI ISL 410540 | P5L | M.javanica
EPI ISL 410539 | P1E | M.javanica
EPI ISL 410538 | P4L | M.javanica
MG772933 | ZC45 | R.sinicus
MG772934 | ZXC21 | R.sinicus
KF294457 | Longquan-140 | R.monoceros
KF569996 | LYRa11 | R.affinis
KU973692 | F46 | R.pusillus
KP886809 | YNLF 34C | R.ferrumequinum
KP886808 | YNLF 31C | R.ferrumequinum
NC 014470.1 | BM48-31 | R.blasii
KY352407 | BtKY72 | Rhinolophus sp.
0.1

**Figure S1. Uncollapsed maximum likelihood phylogenetic tree of the full-length *Sarbecovirus* sequences (Related to Figure 1)**

The full-length sequences (~30,000 bp) of SARS-CoV-2 (Wuhan-Hu-1 as a representative), SARS-CoV-2-related viruses from bats (n=4) and pangolins (n=4), SARS-CoV (n=190), SARS-CoV-related viruses from civets (n=3) and bats (n=54), and outgroup viruses (n=2; BM48-31 and BtKY72) were analyzed. Accession number, strain name, and host of each virus are indicated for each branch. The collapsed tree is shown in **Figure 1A**, and the sequences used are summarized in **Table S1**. The red numbers on the nodes indicates the bootstrap values (>85%). A scale bar indicates 0.1 nucleotide substitutions per site.

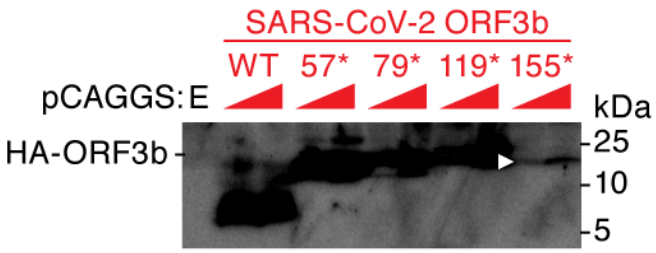


**Figure S2. Visualization of the band of the 155* mutant (Related to Figure 3)**
To visualize the band of the 155* mutant in the Western blotting shown in **Figure 3C**, a highly exposed blot is shown. The band of the 155* mutant is indicated by a white arrowhead. E, empty vector. kDa, kilodalton.
